# Optical Properties of Gelatin Methacrylate and Implications for In Situ Cross-Linking in Spinal Cord Lesions and Engineered Hydrogel Systems

**DOI:** 10.1101/2025.09.15.676407

**Authors:** P. R. Juralowicz, T. J. Bennet, I. Liubchak, Z. Xie, Y. Oloumi Yazdi, T. M. Caffrey, K. C. Cheung

## Abstract

Understanding how light interacts with biomaterials is critical not only for therapeutic delivery in vivo but also for enabling emerging light-based fabrication methods such as volumetric bioprinting. In particular, optical characterization of photosensitive hydrogels like Gelatin Methacrylate (GelMA) provides foundational data for additive manufacturing of complex, tissue-engineered constructs with spatially controlled architecture. GelMA is a promising candidate for use as an injectable hydrogel for spinal cord injury (SCI) repair. This is primarily due to its highly tunable properties, allowing it to mimic tissue. Additionally, GelMA can be optically crosslinked. Optical simulations of this material within the spinal cord can inform whether the cross-linking of this material is feasible and uniform within a particular injury with a chosen exposure paradigm. These simulations require the optical properties of the tissues and materials of interest. However, the optical properties of GelMA have not been studied. The optical properties of GelMA were measured using a double integrating sphere setup and modelled with inverse adding doubling. We measured the absorption coefficient (µ_a_ [mm^-1^]), reduced scattering coefficient (µ_s_’[mm^-1^]), and scattering anisotropy (g [unitless]) for GelMA hydrogels, and examined the effect on these properties when magnetically alignable microstructures were suspended in the hydrogel. We conducted Monte Carlo simulations using these optical properties to determine whether the GelMA hydrogels would be cross-linked under different illumination conditions within a representative spinal cord injury geometry. The depth of hydrogel cross-linking for this representative spinal cord injury geometry was experimentally validated. Overall, it was found that GelMA without rods has an µ_a_ = 0.516 ± 0.025 mm^-1^, with insignificant scattering at 450 nm, GelMA with rods has an µ_a_ = 0.588 ± 0.018 mm^-1^, µ_s_’ = 0.196 ± 0.024 mm^-1^, and g = 0.906 ± 0.014 at 450 nm. Additionally, it was observed that in fluid-filled lesions, external illumination is often sufficient to achieve uniform gel curing. In contrast, fibrotic lesions require an intraspinal approach to adequately deliver light throughout the lesion, although the dose distribution will be nonuniform. Finally, experimental results confirmed the simulation findings, showing that when exposing GelMA to 63.29 mW/cm^2^ of 450 nm light, curing is uniform within the first 4 mm of gel and worsens with depth. These findings support the conclusion that larger volumes of photosensitive GelMA can be feasibly cured within the porcine spinal cord, but an intraspinal exposure will likely be required within human spinal tissue. These findings not only support the feasibility of in situ photopolymerization for SCI repair, but also provide critical optical parameters and validation approaches relevant to bioprinting systems.

## Introduction

Every year, worldwide, an estimated 250,000 to 500,000 people suffer a spinal cord injury (SCI), leading to significant personal suffering and societal costs (1). Despite this, current medical methods only prevent further injury or mitigate symptoms (1). No current treatments attempt to fully reverse the damage. Recent attempts in both *in vitro* and *in vivo* have explored injectable gel-based approaches to provide three-dimensional support for neurite extension within the injury site (2–5). Injectable, space filling hydrogels have been chosen due to how SCI typically manifests in humans. After an acute injury, a fluid filled lesion forms within the spinal cord, and is surrounded by a glial scar (1). This inhibitory environment disrupts signal propagation down the injured spinal cord (1). Hydrogels such as gelatin methacrylate have been shown to promote neuronal growth by filling the fluid filled cavity with a structured scaffolding, while the inclusion of alignable rods within the hydrogel have been shown to guide neurite growth to a desired direction, such as down the length of the cord (5–20). This potential combination of materials could form a minimally invasive treatment for SCI in which a gel is injected into the lesion site and cross-linked. This gel could then act as a scaffolding for neurites to grow through the lesion site.

Injectable scaffolds have emerged as a promising strategy in SCI repair (2,3,5,20–23). These hydrogel systems involve the delivery of a precursor into the lesion site, followed by *in situ* gelation to form a growth conducive matrix. Hydrogel formulations vary significantly in their base material, bioactive additives, and gelation mechanism. Gelation methods can be classified into two categories: physical / reversible hydrogels, and chemical / permanent hydrogels (24). This work focuses on the permanent cross-linking of hydrogels using photo-chemical reaction. Unlike other permanent cross-linking methods, such as small-molecule, polymer-polymer, and enzymatic cross-linking, photo-chemical reactions offer superior spatiotemporal control (24). This differs from other methods as they proceed to completion one initiated. The photochemical approach not only enables precise control over the initiation and termination of gelation but also permits *in situ* modulation of hydrogel mechanical properties, meaning gel properties can be modified by both the formulation as well as the exposure parameters used during administration.

The aforementioned properties offer significant advantages for surgical applications. Firstly, temporal constraints are mitigated. In the previously discussed two-component systems, gelation is initiated upon mixing of gel precursors, necessitating immediate injection to prevent premature gelation. Varying concentration of gel components can extend gelation kinetics, allowing more time for surgical application; however, these modifications risk compromising structural integrity due to potential precursor leakage from the lesion site prior to complete gelation. Photochemical crosslinking can circumvent these issues as gelation is initiated by light exposure, and cross-linking can be completed within minutes (24). Secondly, the ability to modulate hydrogel properties through precise control of the light exposure parameters (time and energy) enables *in situ* modification of material characteristics while ensuring a complete cross-linking reaction.

While photosensitive hydrogels offer significant advantages, their implementation is not without challenges, particular in the method of light delivery. A primary concern is the attenuative properties of biological tissue, particularly in the ultraviolet (UV) and near-UV range, especially as the absorption peak of many photo-initiators reside within this range. (25). Light propagation through turbid media, such as tissue, is governed by absorption and scattering phenomena (26). Biological tissue is highly scattering, causing most light which travels through it to diffusely reflect and exit the tissue surface. This is particularly important for SCI as not only is the spinal cord highly attenuating (27,28), but the injury site is typically encapsulated by spared tissue (1). This spared tissue is comprised of white mater, grey mater, dura matter, and fibrotic scaring, all of which attenuate light as it attempts to travel to the lesion site. These factors require careful consideration of illumination strategies for photochemical gelation systems. External illumination, while the most straightforward approach, requires light to propagate through multiple tissue layers, attenuating significantly before reaching the gel. Internal illumination via an inserted optical catheter could mitigate these issues but may affect cross-link uniformity due to excess exposure at the fiber tip. Computational modelling of the absorption within the spine can shed some light on the final properties of the gel. Optimal exposure paradigms can be chosen based on the absorption model determined by these simulations, with the best paradigm having the most uniform absorption within the gel.

Optical simulations allow researchers to model how light energy is absorbed in tissue (28,29). These Monte Carlo simulations simulate the path of an individual photon of light as it travels through the tissue. This individual simulation is then repeated millions of times to determine the absorption distribution (30). Monte Carlo simulations require the user to input both the optical properties and geometry of each tissue of interest. Most common human tissues have their optical properties published. Despite the wealth of research into the growth potential of hydrogels, and the usefulness of photosensitive hydrogels, the optical properties of any candidate gel have not been previously reported. These simulations can also apply to 3D bioprinting with GelMA (31).

The propagation of light through tissue is often described by the radiative transport theory (25). Each tissue is modelled as a homogeneous turbid media that both absorbs and scatters incident light (25). Absorption is the process in which a photon of light being consumed by some molecule, and having its energy converted into some other process (heat, chemical reaction, fluorescence, etc.), while scattering is the process in which a photon’s trajectory is altered via an interaction with a boundary separating different refractive indices (25). In biological media, this is often the boundary between lipids and aqueous components (25). The optical properties of each tissue can be fully characterized by three parameters: The coefficient of absorption, µ_a_ [mm^-1^], which describes the probability of absorption per unit path length; The coefficient of scattering, µ_s_ [mm^-1^], which describes the probability of scattering per unit path length; and the scattering anisotropy, g [dimensionless, between 0 and 1], which describes the angular distribution of photon trajectories after a scattering event (25,30). A fourth derived optical properties can be calculated with g and µ_s_, called the reduced scattering coefficient, µ_s_’[mm^-1^], which simplifies photon transport as a random walk with step size 1/ µ_s_’ with a uniform distribution of post-scattering angles (25).

This work presents the optical properties of our hydrogel formulation, as well as simulated distribution of dosage among several different lesion geometries and exposure positions. With these optical properties and simulations, this works suggests an optimal surgical path forward for future large animal studies, as well as potential clinical studies. The ability to model and experimentally verify spatially patterned light penetration in GelMA also lays important groundwork for designing 3D light fields to fabricate thick, heterogeneous tissue constructs with high spatial fidelity.

## Methods

### Sample Preparation

Our hydrogel precursor contains three components. The gel base, 50% degree of functionalization gelatin methacrylate (PhotoGel50%DS, Cellink), the photo initiator containing tris(2,2’-bipyridyl) ruthenium(II) (henceforth referred to as Ru) and sodium persulfate (SPS) (SIGMA), and magnetically alignable micro-rods to provide structure and enable directed growth (32). The polycaprolactone (PCL) micro rods contain super paramagnetic iron oxide particles (SPIONs) to enable magnetic alignment as well as the fluorophore Nile red to enable imaging by fluorescence microscopy. Each rod is approximately 5 µm wide, 5 µm tall, and 50 µm long. First, a stock of each component is created. A 15% w/v stock of GelMA is created by adding 3.35 ml of PBS to 500 mg of freeze dried GelMA. This is mixed over night at 45°C with a magnetic stir bar at 200 rpm. It is essential that this stock is protected from ambient light (i.e. wrapped in aluminum foil). Additionally, a 10 mg/ml Ru stock and a separate 10 mg/ml SPS stock are also created. The Ru stock can be created and stored in a 3°C fridge, but the SPS stock must be made fresh. A combined Ru + SPS stock cannot be created as the resulting solution is unstable (33). Creation of a stock of alignable rods depends on the source. In this work, we received a stock of rods from a collaborator (16). Once all stocks are prepared, a solution of 6% w/v GelMA with 0.2 mM Ru and 2.0 mM SPS is created. In cases where rods are experimentally required, the stock is added to achieve a 1% v/v concentration. This concentration is chosen as previous work has shown that to be the best for neuronal growth (32). Once the gel solution is complete, samples are prepared (figure 1). GelMA formulations, and their curing method, are shown in Tables 1 and 2. For aligned samples with rods, alignment is achieved with two stacks of two 38.1 mm diameter, 19.05 mm thick N52 permanent neodymium magnets (K&J Magnetics, Inc.) placed in a parallel configuration (field strength of 42 mT in centre of the magnetic configuration raised 20 mm from the base of the magnets, 7.2 cm magnet centre-to-centre distance).

**Figure 1:**
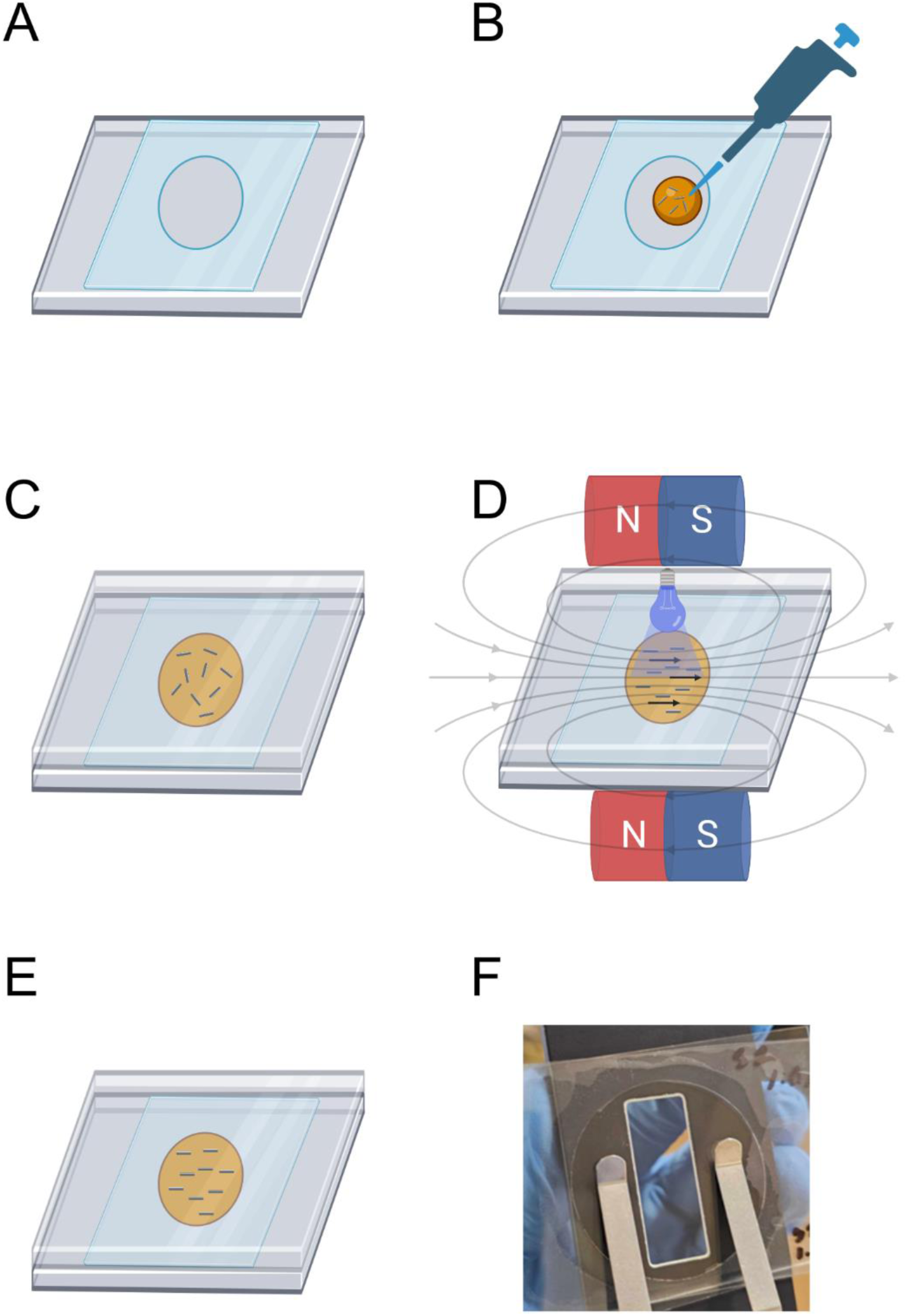
Sample preparation for optical characterization experiments. A) A rectangular section of 0.1 mm thick transparency film (Type E, Cannon) is sectioned. A sample hole is made using a 38 mm diameter punch. The hollowed film is placed on top of a 1mm thick glass slide (Fisherbrand microscope slides). The glass slide thickness is confirmed with a micrometer. B) 250 microliters of prepared GelMA formulation (see table 1 for a list of formulations) is pipetted into the hollow transparency film. C) The top glass slide is placed on top, slowly spreading the GelMA across the hollowed circle, ensuring no air bubbles are formed. The thickness of this glass slide is also confirmed with a micrometer. D) If rods are present, they are aligned with two magnets. The magnets are kept in place during cross-linking (63.8 mW / cm^2^ 455 nm spotlight for 60 s). E) Once cross-linked, the total sample thickness is measured with a micrometer to determine the GelMA thickness (∼0.15 mm). Accurate GelMA thickness is essential for accurate optical properties. F) Example of a prepared GelMA sample, mounted for optical measurements. Created in part with BioRender.com.

**Table 1:** GelMA formulations and curing methods.

| **Code** | **Formulation** | **Curing Method** | **Rationale** |
| --- | --- | --- | --- |
| I | 6% w/v GelMA, 0.2 mM Ru, 2.0 mM SPS | Thermal | Determine the optical properties of GelMA with photo-initiator. |
| II | 6% w/v GelMA, 0.2 mM Ru, 2.0 mM SPS | Photo-crosslinked | Determine if the photo cross-linking process itself affects optical properties |
| III | 6% w/v GelMA, 0.2 mM Ru, 2.0 mM SPS, 1% v/v PCL 5x5x50 μm Rods | Photo-crosslinked | Determine the optical properties of GelMA with photo-initiator and with un-aligned rods. |
| IV | 6% w/v GelMA, 0.2 mM Ru, 2.0 mM SPS, 1% v/v PCL 5x5x50 μm Rods | Photo-crosslinked, with magnet | Determine the optical properties of GelMA with photo-initiator and with un-aligned rods. |
| V | 6% w/v GelMA | Thermal | Determine the optical properties of GelMA without any additives. |

**Table 2:** GelMA curing method descriptions.

| **Method** | **Description** |
| --- | --- |
| Thermal | The sample is placed in a 3°C refrigerator for 15 minutes. |
| Photo-crosslinked | The sample is exposed with 60 mW/cm<sup>2</sup> 455 nm light for 60s. |
| Photo-crosslinked, with magnet | The sample is placed within a 42 mT magnetic field for 60s to align rods within the gel precursor. Then, while remaining in the field, the sample is exposed with 60 mW/cm<sup>2</sup> 455 nm light for 60s. |

### Optical Property Measurement

The optical properties µ_a_ and µ_s_’ were determined using an integrating sphere setup (Cary 7000, Agilent), which can measure the total transmission as well as the diffuse reflectance of our samples. The refractive index of GelMA was measured in a refractometer with white light centered at 589.3 nm at a sample temperature of 23°C (ABBE-3L Refractometer, Spectronic Instruments). The total transmission, diffuse reflectance, and refractive index measurements can be used to calculate the optical properties µ_a_ and µ_s_’. This technique is explained in detail elsewhere (34). Sample size should be modified to best fit the integrating sphere used. This work had an integrating sphere with a 36 mm exit port, requiring a 38 mm sample to fully cover the port. This data was measured across the visible spectrum, from 350 nm to 600 nm. Ballistic transmission, used to measure g, was measured at across the same spectrum using two aligned polarizing filters to reject scattered light (Cary 7000, Agilent). This data was then applied to the inverse adding doubling (IAD) algorithm, which then estimated the desired optical properties. An open source software implementation (OMLC Inverse Adding Doubling Software) in addition to a custom python script that reformatted raw data was used to automatically perform the calculations (35,36). To further improve accuracy, the software was provided the refractive indices of the glass slides as well as the gel across the visible spectrum. The refractive index spectrum of soda lime glass was from (37), while the refractive index spectrum of GelMA was based on the spectrum for water, scaled such that the refractive index at 589.3 nm was identical to the value measured experimentally (38). Analysis of the optical properties of each sample was completed in GraphPad Prism, with t-tests used to compare pairs of formulations, and a 95% confidence interval constructed using the sample standard deviation, mean, and t-distribution.

### Monte Carlo Simulations

Monte Carlo simulations were performed using the experimentally derived optical properties of GelMA as well as literature values for the white matter, grey matter, dura, and cerebral spinal fluid (39–48). Importantly, simulations only used the optical properties of GelMA with rods, as this most closely mimics the *in vivo* application. Mcyxz, a compiled C++ program with a MATLAB interface, was used to compute the simulation (49). Tissue geometries were based on histological image of an injured rat spinal cords, which were scaled up to the approximate size of a porcine spinal cord (Table 5). Two sample geometries were used for this study, one with a fluid filled lesion, and one with a fibrotic lesion. It is assumed that the GelMA injection does not disturb the geometry of the tissue or the debris and uniformly fills the entire lesion site. The MATLAB interface script is used to generate the simulation parameters used by mcxyz. This script scales the spinal cord image, adds the dura and CSF, sets the illumination angle wavelength of the light, and the optical properties of each tissue type. Optical properties for the gel are derived from formulation III. Since spinal cord geometry was converted from an image, scaling was achieved by setting the distance per pixel in a MATLAB interfacing script such that the simulation geometry matched the average size of a pig spinal cord (50). See *figure 2* for details. For each geometry, three exposure paradigms were used: An external illumination with a uniform 4 mm diameter beam, a subdural illumination with a uniform 4 mm diameter beam, and a fully internal isotropic point source, meant to represent an ideal case where a uniformly emitting fiber optic is placed near the center of the lesion. Note: simulation results will be displayed in units of 1/cm^3^, meaning the results are agnostic of exposure power.

**Figure 2:**
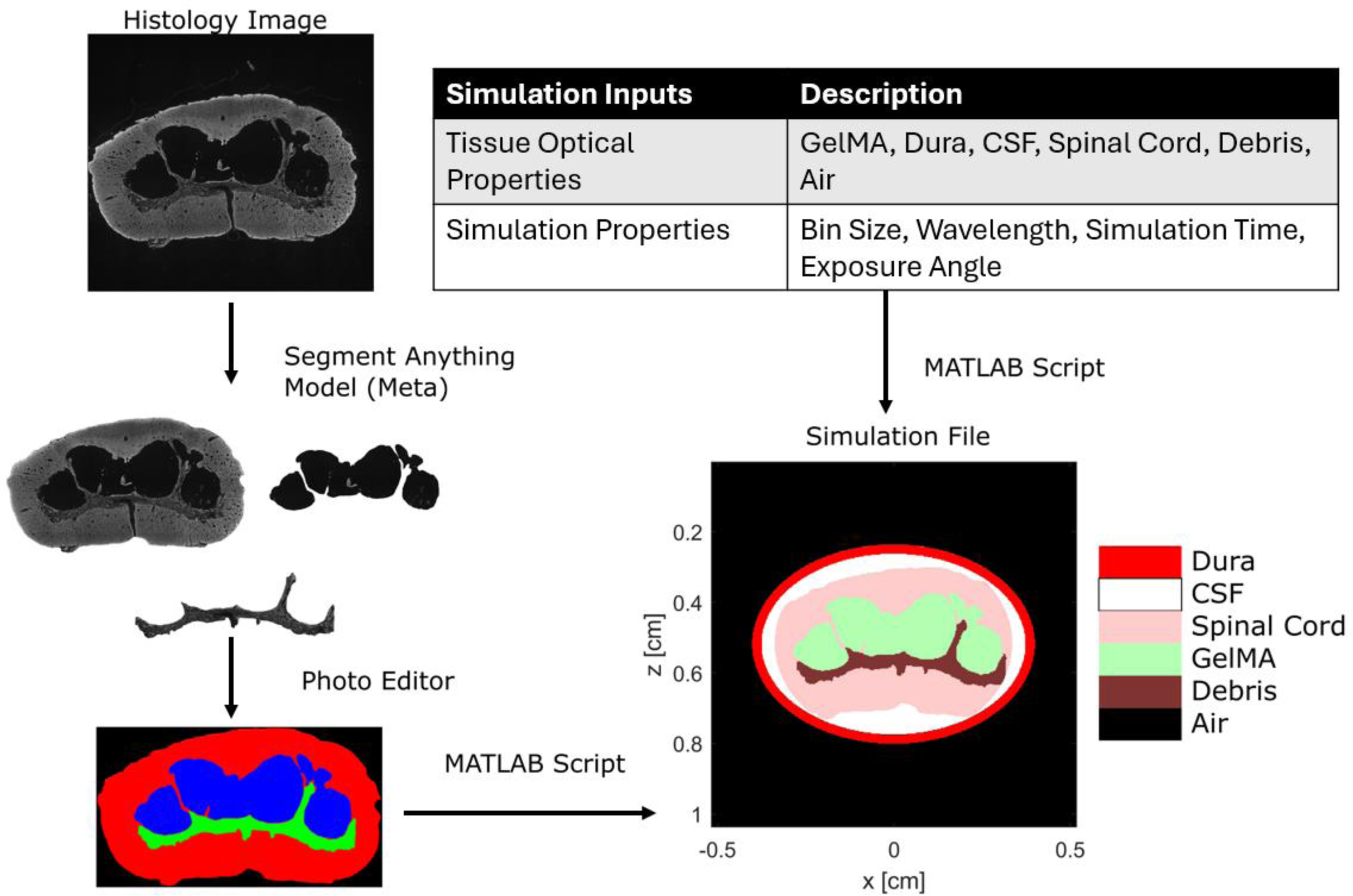
Simulation Setup. A histological image of the spinal cord, with a lesion, is collected. This image is then fed into a segmentation model (Segment Anything, Meta AI) to separate the lesion site (blue), spinal cord (red), and debris (green). Finally, this, along with the optical properties of all tissues of interest are provided to a MATLAB script which generates the simulation file. Within the script that converts the segmented image into the final simulation geometry, the segmented lesion is assumed to be uniformly filled with GelMA. A bin size of 0.0264 mm was used.

**Table 3:** Tissue optical properties from literature.

| Tissue | $\mu_a$ [mm<sup>-1</sup>] | $\mu_s'$ [mm<sup>-1</sup>] | g |
| --- | --- | --- | --- |
| Dura (39–41) | 0.08 | 5.45 | 0.80 |
| CSF (42) | 0.00003 | 0.01 | 0.16 |
| Spinal Cord (39,43–46) | 1.10 | 3.42 | 0.90 |
| Debris (39,43–46) | 2.11 | 3.42 | 0.87 |
| Air | 0.00001 | 0.0 | 1.00 |

**Table 4:** GelMA optical properties at 450 nm, 95% confidence interval.

| Formulation | $\mu_a$ [mm<sup>-1</sup>] | $\mu_s'$ [mm<sup>-1</sup>] | g |
| --- | --- | --- | --- |
| 6% w/v GelMA, 0.2 mM Ru, 2.0 mM SPS, Thermal (I) | 0.527 ± 0.027 | N/A | N/A |
| 6% w/v GelMA, 0.2 mM Ru, 2.0 mM SPS, Photo-Crosslinked (II) | 0.516 ± 0.025 | N/A | N/A |
| 6% w/v GelMA, 0.2 mM Ru, 2.0 mM SPS, 1% v/v PCL<br>5x5x50 $\mu\text{m}$ Rods, Photo-Crosslinked, Unaligned (III) | $0.588 \pm 0.018$ | $0.196 \pm 0.024$ | $0.906 \pm 0.014$ |
| 6% w/v GelMA, 0.2 mM Ru, 2.0 mM SPS, 1% v/v PCL<br>5x5x50 $\mu\text{m}$ Rods, Photo-Crosslinked, Aligned (IV) | $0.576 \pm 0.086$ | $0.233 \pm 0.054$ | $0.889 \pm 0.021$ |
| 6% w/v GelMA, Thermal (V) | $0.035 \pm 0.0005$ | N/A | N/A |
| Refractive index of 6% w/v GelMA at 23°C | 1.34 (does not change between formulations) |  |  |

**Table 5:** Rat and porcine spinal cord geometries, lumbar spinal cord (50).

| Animal | Height | Width |
| --- | --- | --- |
| Rat | 2-2.5 mm | 2.5-4 mm |
| **Pig** | 5-6 mm | 7-10 mm |
| **Human** | 7-8 mm | 9-10 mm |

Multiply the results by the total exposure power to get the power volume density.

### Photopolymerization crosslink depth

*Figure 3* describes the process of creating a validation experiment for the measured optical properties. A column of GelMA is created using a PDMS mold. This column is then cross-linked and sectioned into 2-mm disks, and the sol fraction of each disk is measured to determine the degree of gel cross-linking as a function of distance from the light source. Sol fraction refers to the fraction of polymer that remains outside of the crosslink network (51). It is governed by the following equation:

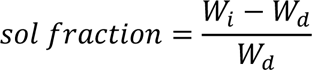

**Figure 3:**
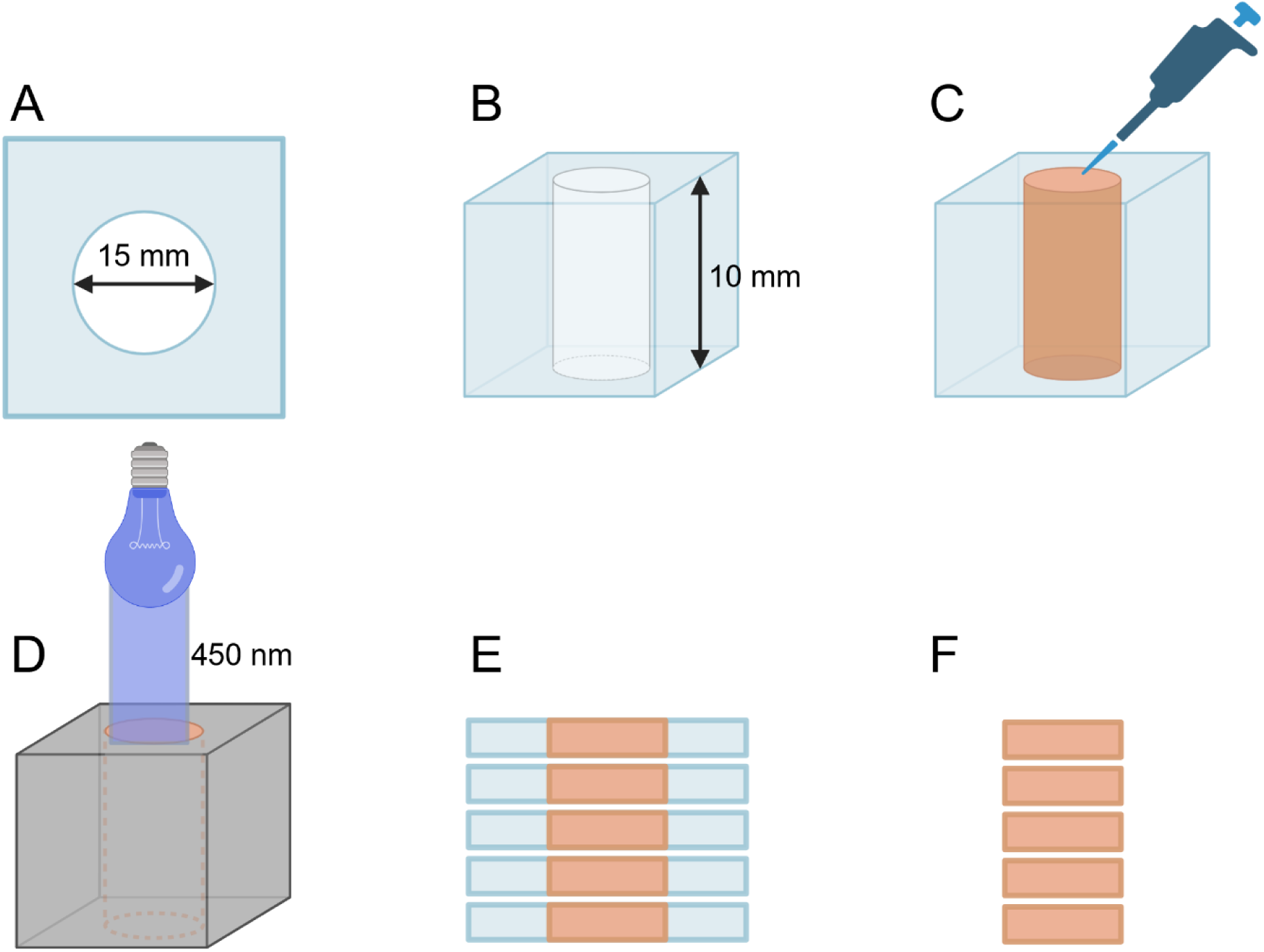
Experimental setup to determine photopolymerization cross-link depth. A) Square sheets of 2 mm thick PDMS with a 15 mm cylindrical hole are cast in a 3D printed mold. B) These sheets are then plasma-bonded together, ensuring alignment of the hole. This forms a PDMS mold for a column, with a total height of 1 cm. C) The mold is filled with a GelMA formulation while on a heat block to prevent thermal gelation. Before light exposure, the hydrogel column is thermally gelled for 20 minutes at 4°C to enhance structural stability. D) An opaque 3D printed PLA shroud is placed over the pillar to ensure the light must first pass through the GelMA. The GelMA column is exposed with 450 nm light with power density 63.29 mW/cm^2^ for 60 s. The column is thermally crosslinked for an additional 5 minutes after exposure. E) PDMS sheets and the embedded gel column are sectioned using a razor, creating hydrogel disks. F) GelMA disks are removed, and the sol fraction of each GelMA disk is measured. Created in part with BioRender.com

Within this equation are *W*_*i*_, the initial dry weight after cross linking, and *W*_*d*_, the dry weight after all un-crosslinked material is removed (51). *W*_*i*_ is measured by taking the sample after photo-crosslinking and freeze drying. This is done to accurately measure the weight of the gel components (both crosslinked and not) without water. *W*_*d*_ is measured after incubating the sample in 2 mL 37*°C* PBS for 24 hours and a second freeze-drying. This is done to accurately measure the weight of only the successfully crosslinked gel components. A lower sol fraction indicates more complete crosslinking. This experiment is completed with formulation II and IV (**Table 1).** The gel within the column is thermally gelled before exposure. Without this, cured gel section would sink into the uncured sections. Note that thermal gelation is reversable by heating. Finally, data was analyzed using one-way anova.

## Results

### Optical Properties

The optical properties of all formulations were measured from 350 nm to 750 nm in 5 nm increments. Light absorption peaks at 450 nm (the absorbance peak of Ru) in all formulations in which photoinitiator is present. There is an additional absorbance peak in all formulations in the ultraviolet (UV) range. This peak is the only peak present in formulation V, implying that pure GelMA absorbs within the UV range. This, in addition to Ru’s UV absorption, leads to significant UV absorption in formulations I-V. Scattering was measurable in formulations with a scattering agent (rods), and uniform across the visible range. Paired t-tests and ANOVA confirmed significant differences in absorbance between rodless and rod containing formulations, significant difference in absorbance when a photoinitiator is included in a formulation, no significant difference between thermal and photo-crosslinking, and no significant difference between aligned and unaligned rods.

### Monte Carlo Simulations

The following two geometries were used for all Monte Carlo simulations. Both were based on rat histological images and scaled to the size of a porcine spinal cord. The porcine spinal cord was chosen as a porcine model of spinal cord injury is the next step for this treatment before clinical trials. The first geometry is a primarily fluid filled lesion (*Figure 5 A*). The second is a fibrotic lesion with debris *(Figure 5 B).* It was assumed that the injection would fill the lesion uniformly and not displace any debris within the lesion itself. Note: simulation results will be displayed in units of 1/cm^3^, meaning the results are agnostic of exposure power. Multiply the results by the total exposure power to get the power density. The spinal cord was represented as one tissue type based on the optical properties of white matter; no distinction is made between white and grey matter. The reasoning behind this is as follows: First, it was difficult to separate white and grey matter in the histology images. Secondly, the formation of the lesion site destroys much of the grey matter (1). Finally, white matter optical properties are similar, but slightly more attenuating, than the optical properties of grey matter (combining the absorbance and scattering coefficient yields an effective 3.83 mm^-1^ attenuation coefficient for white matter, 3.02 mm^-1^ for grey (28)), making the simulation the lower-bound of energy deposition within the lesion site.

**Figure 4:**
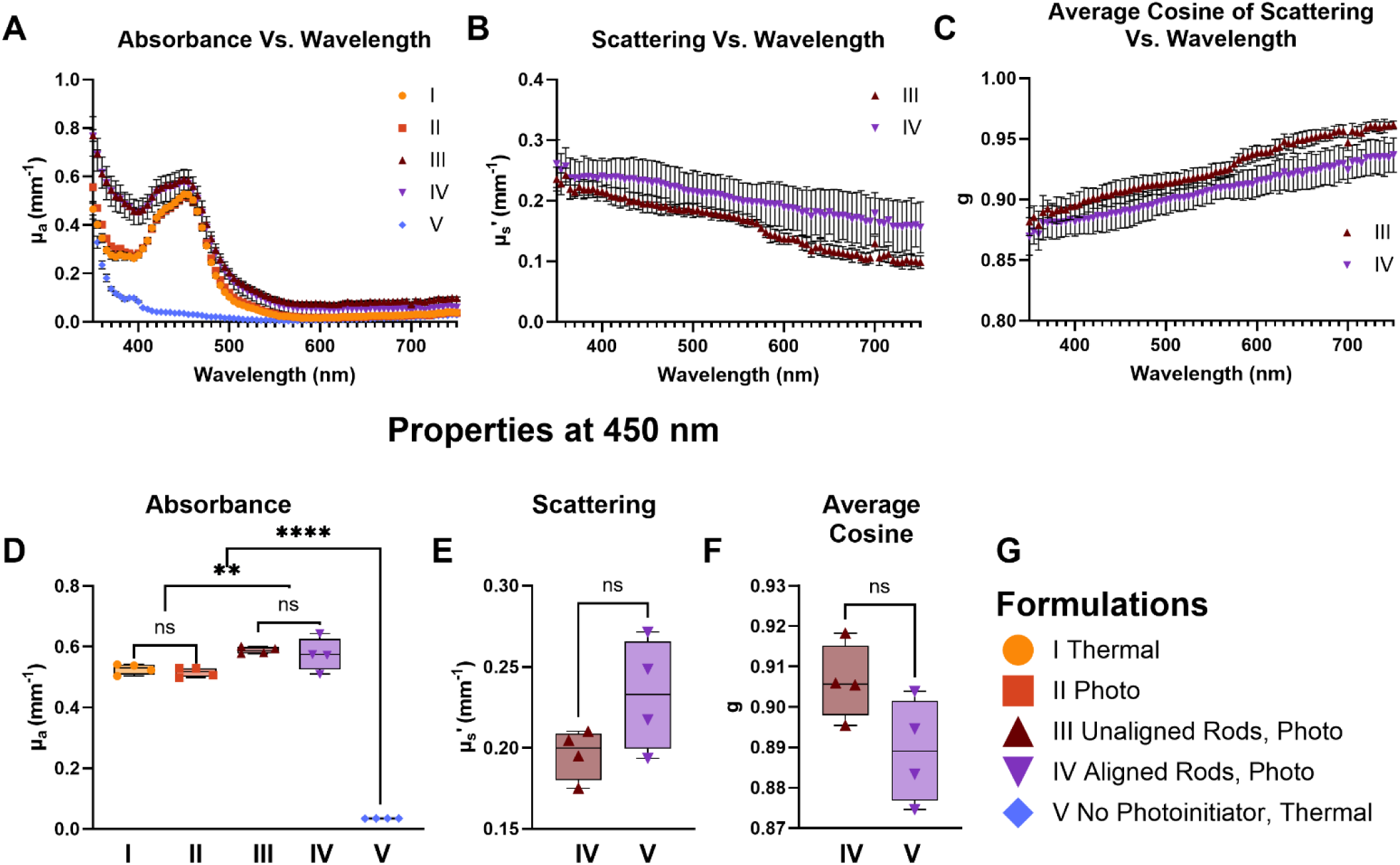
Optical Properties (n=4). All formulations use 6% w/v GelMA with a photoinitiator of 0.2 mM Ru and 2.0 mM SPS as base, unless otherwise noted. A-C) Optical property spectrums. Note: GelMA without rods did not scatter significantly, so no scattering properties were determined. D-F) Optical properties at 450 nm, statistically significant differences noted (** p ≤ 0.01, **** p ≤ 0.0001, ns p > 0.05). G) Formulation Legend

**Figure 5:**
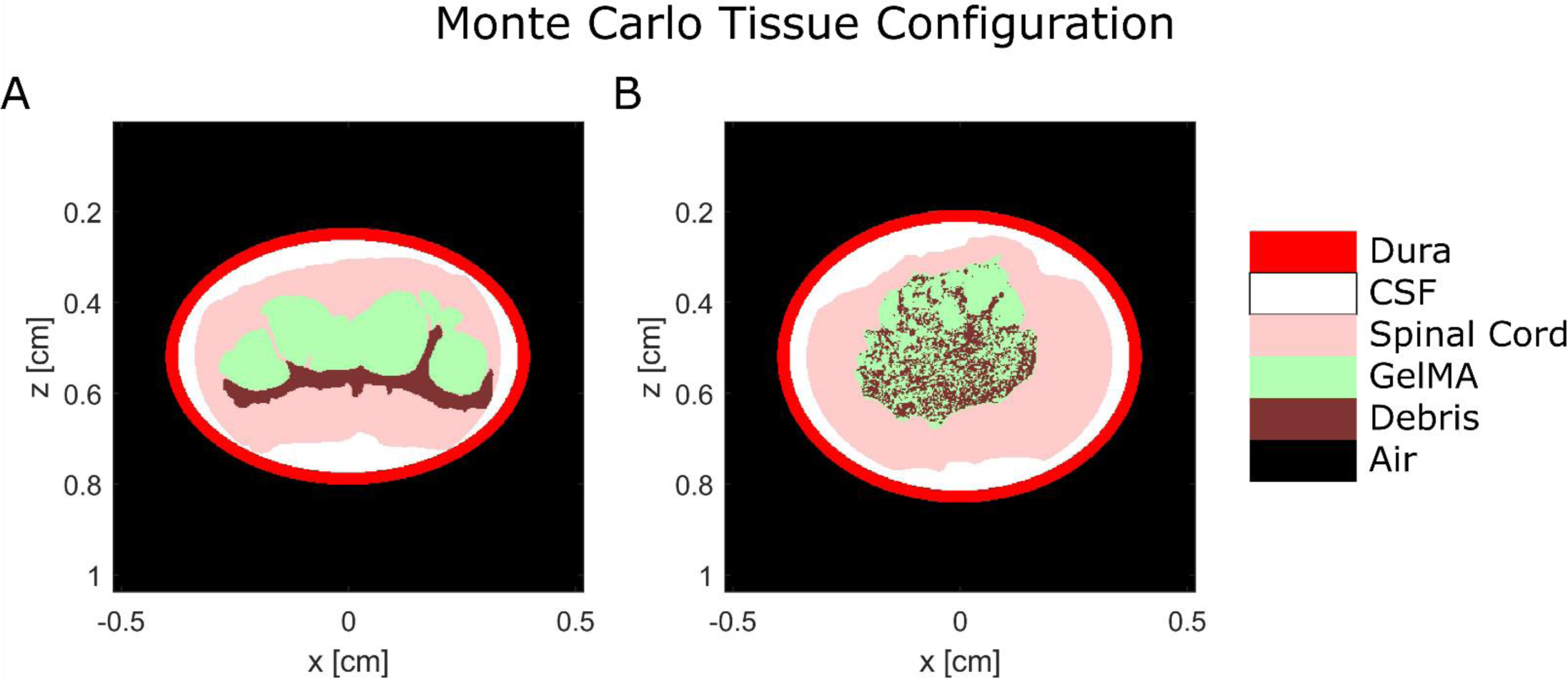
Tissue geometries for the Monte Carlo simulations. A) an initially fluid filled lesion. B) an initially fibrotic lesion. Note: these simulations assume that when the GelMA is injected into the lesion, it replaces any fluid within the lesion site but does not disturb the tissue nor debris.

*Figure 6* displays the results of six Monte Carlo Simulations within the two geometries. Each geometry had three separate exposure paradigms. The first paradigm, the external exposure, placed a uniform, 4 mm wide spotlight above the dura. The second, the subdural exposure, places the same spotlight just underneath the dura, within the arachnoid space. The third, the intraspinal exposure, places a point source in the center of the lesion.

**Figure 6:**
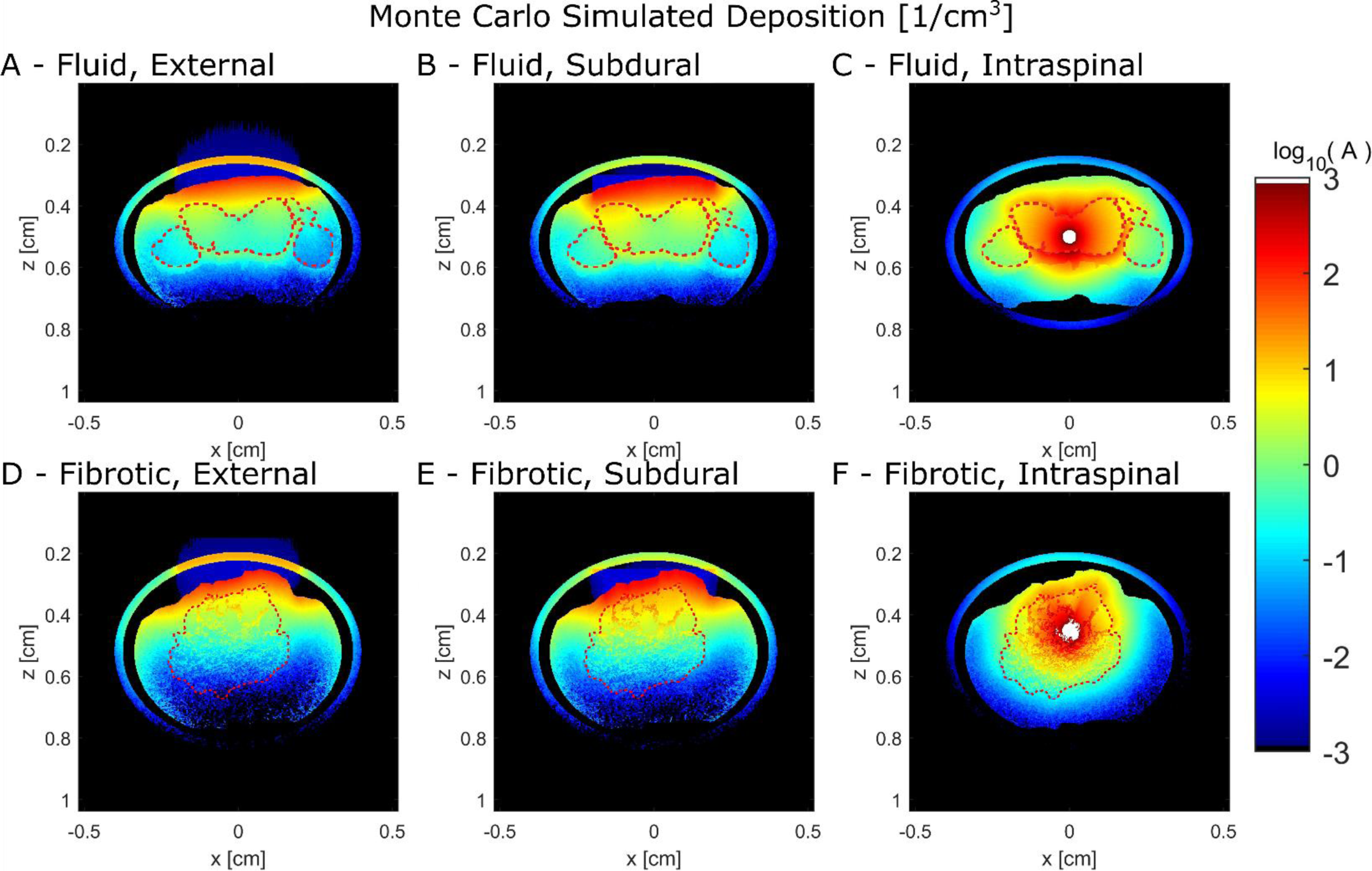
Monte Carlo simulation deposition results (1/cm^3^, or alternatively W/cm^3^/W delivered). The lesion site is marked by a dashed red line. Any deposition above 10^3^ mW/cm^3^ is white, for ease of illustration. A, B, C) Results for a fluid filled lesion from an external, subdural, and intraspinal exposure, respectively. D, E, F) Results for a fibrotic lesion from an external, subdural, and intraspinal exposure, respectively.

*Figures 7* and *8* show an alternative display of the results from Figure 6. Each section shows the deposition long the dashed line shown in the heat map, which runs along the center of the lesion. This shows the uniformity of the dose along the lesion depth. A uniform dose will ensure uniform final mechanical properties, which encourages uniform neurite growth.

**Figure 7:**
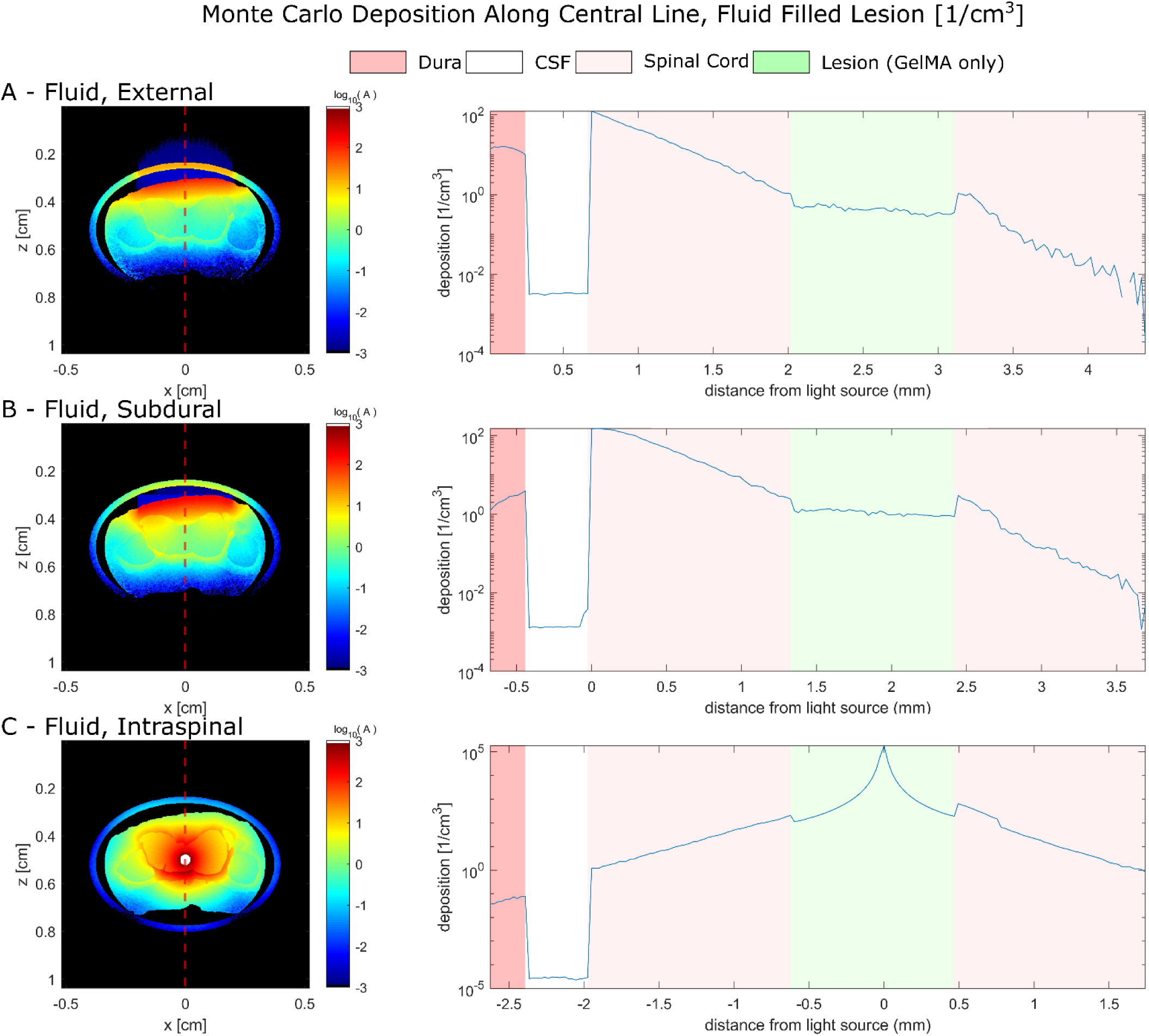
Monte Carlo simulation results for a fluid filled lesion, graphed along a 1D path through the center of the lesion (red dashed line), logarithmic scale in deposition, tissue regions marked. Any deposition above 10^3^ mW/cm^3^ is white, for ease of illustration. Note: The legend reflects the tissue order along the line, with the left most region being the dura. The lesion in this figure is entirely filled with GelMA and uses GelMA’s optical properties. A) External. B) Subdural. C) Intraspinal.

**Figure 8:**
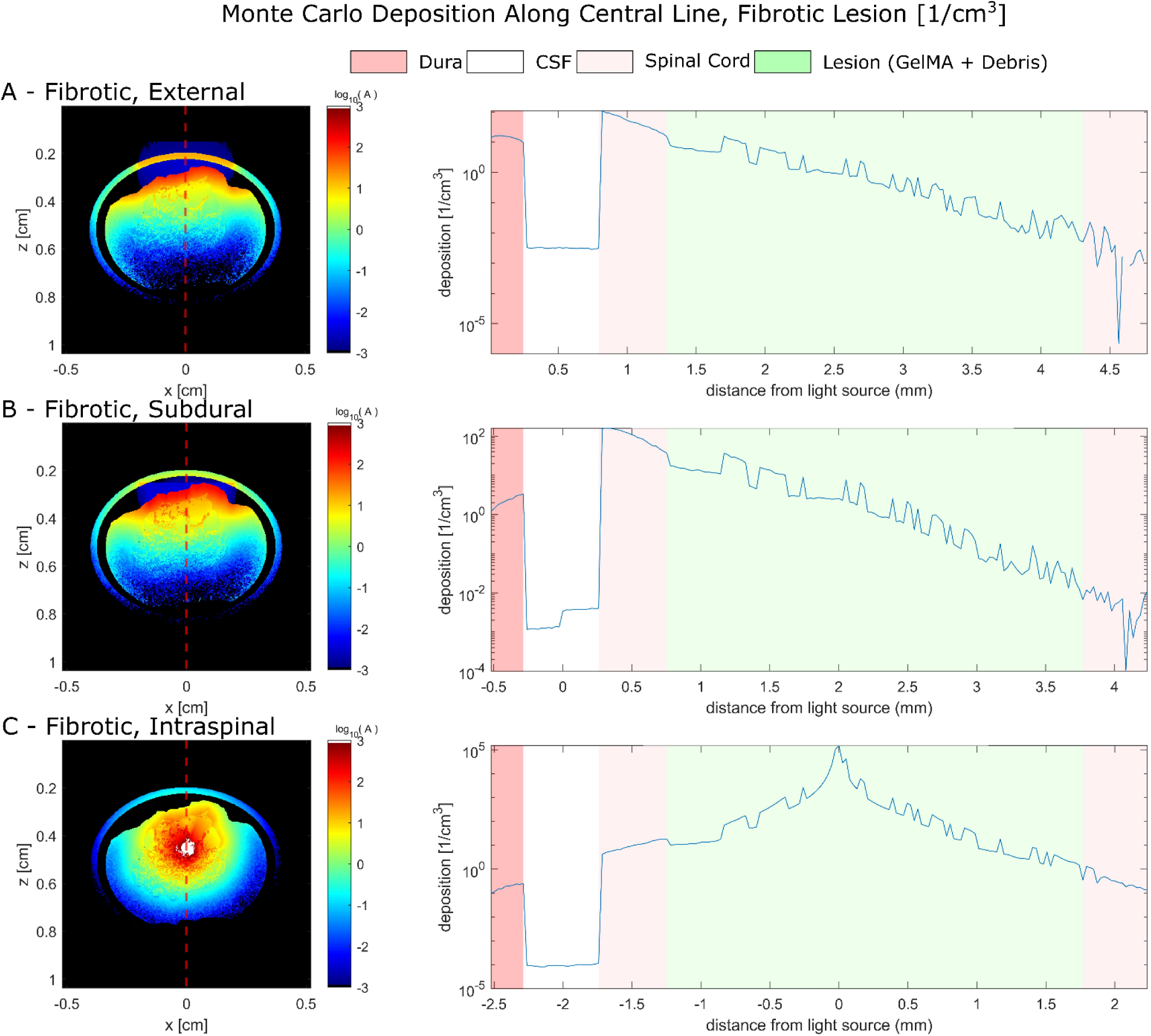
Monte Carlo simulation results for a fibrotic lesion, graphed along a 1D path through the center of the lesion (red dashed line), logarithmic scale in deposition, tissue regions marked. Any deposition above 10^3^ mW/cm^3^ is white, for ease of illustration. Note: The legend reflects the tissue order along the line, with the left most region being the dura. The bumps and valleys within the lesion are due to the presence of debris. The debris has a higher absorption coefficient then the gel, leading to spikes in deposition. Within the lesion, any valley is the gel, while peaks are debris. A) External. B) Subdural. C) Intraspinal.

### Photopolymerization cross-link depth

*Figure 9* describes the experimental and simulated results for photopolymerization cross-link depth experiment. A low sol fraction corresponds to low amount of remaining un-crosslinked polymer, while high sol fraction corresponds to a high amount of remaining un-crosslinked polymer. Experimentally, GelMA curing was consistent for the first 4 mm. Beyond 4 mm, the sol fraction increased, indicating decreased crosslinking efficiency. This is also seen in the simulation results, where beyond 5 mm depth the power quickly drops to less than 10 mW/cm^3^. GelMA containing rods experienced worse curing both experimentally and in simulation, due to both the increased absorption coefficient (due to the Nile Red contained in the rods) as well as the scattering of the rods themselves. This is apparent in *figure 9 C* and *F*, as it is clearly seen that rodless GelMA has a higher penetration depth, and the light path remains primarily ballistic. However, due to the scattering of the rods, an effect known as bunching caused slightly higher deposition within the first slice, leading to a higher sol fraction.

**Figure 9:**
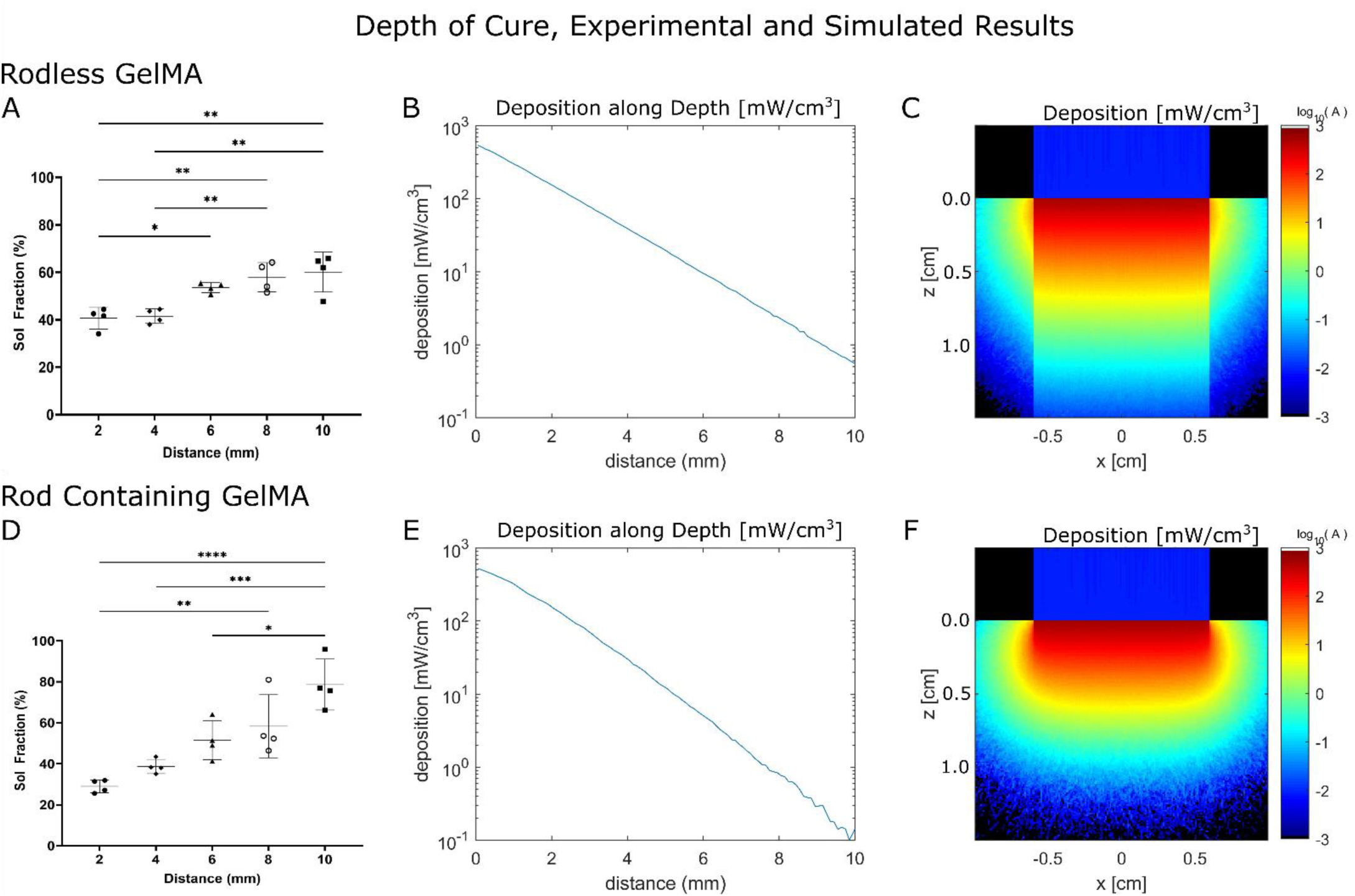
Photopolymerization cross-link depth results for rodless and rod containing GelMA, experimental and simulated. All experiments completed with a 63.8 mW/cm^2^ 450 nm spotlight. A-B) Rodless GelMA. A) Experimentally measured sol fraction. B) Simulated deposition along the center of the column. C) 2D results in an infinite thickness sample of GelMA without rods. D-F) Rod Containing GelMA. D) Experimentally measured sol fraction. E) Simulated deposition along the center of the column. F) 2D results in an infinite thickness sample of GelMA with rods. noted (* p ≤ 0.05 ** p ≤ 0.01, *** p ≤ 0.001 **** p ≤ 0.0001)

## Discussion

### Optical Properties

Absorption reduced scattering, and average cosine of scattering (g) spectra were calculated using IAD (*Figure 4 A-F*). As expected, the absorbance spectrum of GelMA is very similar to the absorbance spectrum of Ru, with a peak around 450nm (*Figure 4 A*) (52). Estimating the absorbance coefficient at 450 nm using the extinction coefficient of Ru and a concentration of 0.2 mM gives an absorbance of 0.67 mm^-1^, compared to an experimentally measured 0.53 mm^-1^ in rodless GelMA. This discrepancy is likely explained by uncertainties in the measurements of Ru extinction coefficient, and uncertainties of various aspects of the samples created for this experiment (Ru concentration, sample thickness, measured transmission and reflectance). Even if individually these uncertainties are small, they can total to the discrepancy seen here. There is a discrepancy between the absorbance spectrum of Ru and Ru + GelMA in the near UV range. This absorbance is caused by the GelMA itself, as evident by the spike in absorbance seen in the pure GelMA formulation. The presence of rods also increases absorbance due Nile Red labeling. Without rods, GelMA scatters insignificantly. IAD could not produce µ_s_’ or g for rodless samples. Both rod containing formulations had uniform and forward scattering across the visible spectrum. The aligned formulation had significantly higher variance compared to the un-aligned, and increased (but not statistically significant) scattering. This could be due to anisotropic rod alignment. This difference of properties can be measured in anisotropic human tissue, such as white matter, where the scattering significantly decreases when light travels along the axon as opposed to perpendicular to it (28).

Finally, *Figure 4 D-F* shows the optical properties of all GelMA formulations at 450 nm. Unpaired t-test were run between thermally and crosslinked GelMA, aligned and unaligned rod containing GelMA, rod and rodless GelMA, and pure GelMA and photoinitiator containing GelMA. There is no significant difference in absorbance at 450 nm between thermally and optically crosslinked GelMA. There is no significant difference in absorbance between aligned and unaligned rod containing GelMA. There is a significant difference in absorbance between rodless and rod containing GelMA, likely due to the presence of Nile Red in the rods. There is a significant difference in absorbance between pure GelMA and GelMA with a photoinitiator, and absorbance at 450 nm is almost entirely dependent on Ru, with minor contributions from any dye in the rods. Unexpectedly, there is no significant difference in the scattering properties of GelMA containing unaligned or aligned rods. This is unexpected as in other turbid media with anisotropic arrangements, such as in white matter, there is significantly different optical properties depending on the photon’s direction of travel (28). In white matter, there is significantly more scattering if the light travels perpendicular to the direction of axonal growth. A similar phenomenon should be seen between the aligned and unaligned rods in principle. This could be mitigated by the sample creation method. Since a droplet of gel is pressed between two glass slides, the flow of gel as it spreads across the glass surface may partially align rods with the sample plane. For a given rod that is out of alignment with the plane of the sample, the flow of the gel as it is squished by the two glass slides will place a torque upon the rod, aligning it with the direction of flow. As the gel is flowing across the sample, the flow direction is parallel with the glass slide and therefore may align rods with the sample plane. This was seen (but not quantified) during microscope imaging of unaligned samples, as although there was a high degree of variance in the angel of the rod within the plane of the sample, no rods seemed to be angled upwards, towards the lens of the microscope. A summary of optical properties can be found in **Table 4**. For 3D bioprinting, the photoinitiator should ideally have peak absorbance beyond 400 nm to avoid interference with the inherent UV absorption of GelMA. Photoinitator concentration should balance layer height with curing time. The Ru concentration in our formulation allows for acceptable penetration depth (especially for the thin layers seen in bioprinting (31)), but requires an extended exposure time. Increased photoinitiator concentration would hasten reaction dynamics while lowering penetration depth, which is irrelevant for small layer sizes.

### Monte Carlo Simulations

*Figure 5* shows the two geometries chosen for the Monte Carlo simulations. The geometry of the spinal cord itself was gathered from histological images taken of rat spinal cords that underwent a contusion injury. As discussed in the methods, these images were then segmented to separate the tissue types of interest (white matter, lesion, debris), and scaled to the size of a pig’s spinal cord. This path was chosen, as opposed to collecting images of a pig spinal cord, as the quality of the histological images was far greater than that of any pig spinal cord images we could receive. To ensure that the scaling process was accurate and did not distort the relative size of the tissues, the scaled cord was compared to CT images of an injured pig spinal cord and adjusted accordingly. The optical properties of GelMA within the simulation were modelled as those of formulation III (6% w/v GelMA with unaligned rods) as unaligned rods make a more spatially homogeneous gel, which better fits the assumption of the simulation. It was also assumed that the injection of GelMA into the lesion would not disturb the lesion nor debris geometry and would completely fill the geometry.

*Figure 6 A-C* shows the results of several Monte Carlo simulation in a fluid filled lesion. In all cases, light can penetrate depth entirely and dose the gel. External and subdural illuminations struggle with exposure on the far sides of the lesion, in particular the left and right-side lobes. As mentioned previously, the highly forward scattering of the gel combined with its low scattering coefficient (relative to the surrounding tissue) means that light is primarily spread by the scattering of other tissue layers as well as reflections along the gel-tissue barrier. This phenomenon is shown in *figure 6 A* and *B*, as the light can spread beyond the initial uniform 4 mm beam and partially dose the side lobes of the lesion. This spreading also ensures more uniform dosage in the lesion body. An intraspinal approach, shown in *figure 6 C*, does enable adequate dosing across the entire lesion, but the dose is incredibly non-uniform. Without the spreading caused by the white matter and dura, the deposition is almost entirely governed by the exponential decay of the light as it is attenuated by the gel.

*Figure 6 D-F* shows results of an additional set of Monte Carlo simulations in a fibrotic lesion. The debris in the lesion significantly attenuate light as it travels through the lesion. This is especially seen in *figure 6 D*, where the bottom of the lesion receives little exposure. In a subdural exposure (*figure 6 E*), the entire lesion is adequately dosed, but the difference in absorption throughout the depth of the lesion would cause nonuniform cross-linking of the gel, harming the therapeutic abilities of the gel. The intraspinal approach (*figure 6F*) also adequately doses the entire lesion but like in the fluid filled lesion (*figure 6 C*), the dosing is extremely non-uniform. The only respite is fluid filled lesion are the most common presentation in humans (1).

*Figures 7* and *8* demonstrate the uniformity of cure by showing deposition of energy along the center of the lesion. Uniform deposition will present as a flat line within the “lesion” region of the graph. Of the attempted lesion geometries and exposure paradigms, only the fluid filled lesion under the external or subdural exposure had uniform deposition along the depth of the gel (*Figure 7 A, B).* Fibrotic lesions under the external or subdural exposure paradigm showed an exponential decrease in deposition (looks linear on the graph, as the y-axis is logarithmic) along the lesion depth, with the external approach failing to cure near the bottom of the lesion (*Figure 8 A, B).* Intraspinal exposures present the worst uniformity, with super exponential decay, which is to say in the graph, the deposition appears to decay exponentially despite the logarithmic scaling of the y-axis (*Figure 7 C, Figure 8 C).* Despite the benefits of coverage and ensure the entirety of the gel is cured, the extreme variance in deposition could lead to extremely nonuniform mechanical properties.

Both exposure geometries emphasize the necessity of pre-operative planning and imaging, and how dosing issues will further increase in human spinal cords. Whether it be the side lobs in the fluid filled lesion or fibrotic blockage, it is probable the light will have trouble reaching across the entire lesion. Proper pre-operative imaging and simulation can direct the location and number of injections / illuminations necessary to ensure proper cross-linking. A sufficiently large fluid filled lesion that is close to the dorsal surface, or an open lesion, could be cross-linked adequately with a large spotlight external illumination.

### Photopolymerization cross-link depth

*Figure 9* describes the experimental and simulated results for photopolymerization cross-link depth experiment. For both GelMA with or without rods, curing is consistent for the first 4 mm and then attenuation increases with depth, affecting crosslinking efficiency. This is an expected result as light power decreases exponentially as it travels through the column. Another unexpected result is how the top two slices of rodless GelMA (*Figure 9 A*) have similar sol fraction despite the deposition decreasing exponentially between them (*Figure 9 B*). This is most likely caused by the resolution of the experimental slices being unable to distinguish between the change in deposition between the two slices. This effect is not seen in rod-containing GelMA, as the higher attenuation enables the resolution of the slices to detect that change in deposition.

Comparing rodless and rod-containing GelMA, the results are as expected. The increased absorption and scattering caused by the inclusion of rods lowers the penetration depth and increases the spread of the light. This is seen in all results. In the experimental results, sol fraction increases quickly with depth (*Figure 9 A, D*). In the simulated deposition along the depth, the linearity of the graph is slightly affected by the spreading of the light, and the overall deposition is less for the same depth in rod containing GelMA (*Figure 9, B, E*). However, the deposition at the top the of column is higher in rod containing GelMA. This is directly seen in the simulated results as well as the experimental sol fraction. The cause of this is a phenomenon called bunching. The scattering of light within a turbid media causes a portion of the photons to scatter so significantly that through multiple scattering events their trajectory reverses, sending them back to the tissue surface. This leads to a disproportionate amount of deposition near the surface. Finally, in the heat map (*Figure 9, C, F*), the depth is smaller in the rod containing GelMA, and the spread of the light is apparent. Unlike the tissue geometry simulations, where the spreading of light caused by the scattering of the tissue layers helps ensure a more uniform dosage, spreading caused by the GelMA itself does not lead to a more uniform cure, it leads to the opposite.

## Conclusion

The optical properties of GelMA were investigated and applied to several biologically relevant simulations as well as validation experiments. Overall, it was found that pure GelMA is optically inert beyond near UV wavelengths. This makes it an excellent base for a variety of photo initiators. When a Ru-based photo-initiator is added, it is an absorbing material, with peak absorption at 450 nm, characteristic of photo-initiator. Without additional additives, GelMA is scatters very little light, especially compared to the surrounding tissue. The addition of alignable rods significantly increases scattering, but GelMA remains less scattering than human tissue. Simulations of GelMA in the injured spinal cord show that large animal dosage is typically possible in fluid filled lesions, with some additional care necessary to ensure proper dosing of especially wide lesions.

However, debris filled lesions present a challenging environment for cross-linking as the debris significantly attenuates the light as it travels through the depth of the lesion. Multiple exposures could mitigate this issue. Finally, simulations were validated through a depth of cross-link experiment, and results were comparable between simulation and experiment. Overall, these results show that GelMA is an excellent candidate hydrogel for SCI repair due to its tunable properties, ease of use, and configurable cross-linking. Beyond applications in spinal cord repair, the characterization of GelMA’s optical properties and corresponding light-dose simulations provide valuable design parameters for emerging volumetric bioprinting techniques. These findings can inform light-based additive manufacturing strategies where uniform and spatially controlled photopolymerization is essential, particularly in the fabrication of complex, tissue-engineered constructs. The ability to predict and validate gel cross-linking in 3D geometries using measured optical properties extends the utility of this work to a range of biomedical applications where precise spatial control over hydrogel curing is critical. However, it should be noted that although GelMA with Ru/SPS has excellent potential within these systems, characterization of alternative photoinitiator systems will be required. Absorption can be estimated using the extinction coefficient of the primary absorber, but scattering tends to require direct measurement. Formulations should ensure candidate photoinitiator systems have peak absorption beyond 400 nm as to not compete with the inherent UV absorption within GelMA itself.

## Acknowledgements

This research has been funded by the Government of Canada’s New Frontiers in Research Fund (NFRF), NFRFT-2020-00238. The Authors thank Garret Frank for his initial guidance, and literature search for the optical properties of the tissue types in the spinal cord. The authors thank Dr. Wolfram Tetzlaff and Dr. Ward Plunet for providing the histological images used in the simulations. The authors thank the Laura De Laporte lab for providing the stock of PCL rods used in this experiment. The authors thank the suggestions and feedback from the entire BioMEMS team. Generative AI (Chat GPT) was used to help edit original non-AI generated text. The following prompt was used: “could you edit this paragraph to sound more professional, such as an excerpt from a scientific journal? ‘[THE ORIGINAL PARAGRAPH].’” The output provided by the AI was then used to inspire edits / vocabulary of the original paragraph. The output was not copied directly into the manuscript. Data used in this research can be acquired upon request from the first author.

## Bibliography

1. Ahuja CS, Wilson JR, Nori S, Kotter MRN, Druschel C, Curt A, et al. Traumatic spinal cord injury. Nat Rev Dis Primers. 2017 Apr 27;3:17018.

2. Yao S, Yang Y, Li C, Yang K, Song X, Li C, et al. Axon-like aligned conductive CNT/GelMA hydrogel fibers combined with electrical stimulation for spinal cord injury recovery. Bioact Mater. 2024 May;35:534–48.

3. Qi Z, Pan S, Yang X, Zhang R, Qin C, Yan H, et al. Injectable Hydrogel Loaded with CDs and FTY720 Combined with Neural Stem Cells for the Treatment of Spinal Cord Injury. Int J Nanomedicine. 2024 May 8;19:4081–101.

4. Velasco-Rodriguez B, Diaz-Vidal T, Rosales-Rivera LC, García-González CA, Alvarez-Lorenzo C, Al-Modlej A, et al. Hybrid methacrylated gelatin and hyaluronic acid hydrogel scaffolds. preparation and systematic characterization for prospective tissue engineering applications. Int J Mol Sci. 2021 Jun 23;22(13).

5. Zhang H, Xu J, Saijilafu. The effects of GelMA hydrogel on nerve repair and regeneration in mice with spinal cord injury. Ann Transl Med. 2021 Jul;9(14):1147.

6. Rose JC, Cámara-Torres M, Rahimi K, Köhler J, Möller M, De Laporte L. Nerve Cells Decide to Orient inside an Injectable Hydrogel with Minimal Structural Guidance. Nano Lett. 2017 Jun 14;17(6):3782–91.

7. Cruz EM, Machado LS, Zamproni LN, Bim LV, Ferreira PS, Pinto LA, et al. A Gelatin Methacrylate-Based Hydrogel as a Potential Bioink for 3D Bioprinting and Neuronal Differentiation. Pharmaceutics. 2023 Feb 13;15(2).

8. de Vasconcelos ACP, Morais RP, Novais GB, da S Barroso S, Menezes LRO, Dos Santos S, et al. In situ photocrosslinkable formulation of nanocomposites based on multi-walled carbon nanotubes and formononetin for potential application in spinal cord injury treatment. Nanomedicine. 2020 Oct;29:102272.

9. Yu H, Fan P, Deng X, Zeng M, Ge L, Xue E, et al. Nerve-Derived Extracellular Matrix Promotes Neural Differentiation of Bone Marrow Stromal Cells and Enhances Interleukin-4 Efficacy for Advanced Nerve Regeneration. Adv Healthc Mater. 2025 Mar;14(7):e2402713.

10. Lim KS, Klotz BJ, Lindberg GCJ, Melchels FPW, Hooper GJ, Malda J, et al. Visible Light Cross-Linking of Gelatin Hydrogels Offers an Enhanced Cell Microenvironment with Improved Light Penetration Depth. Macromol Biosci. 2019 Jun;19(6):e1900098.

11. Fan L, Liu C, Chen X, Zou Y, Zhou Z, Lin C, et al. Directing Induced Pluripotent Stem Cell Derived Neural Stem Cell Fate with a Three-Dimensional Biomimetic Hydrogel for Spinal Cord Injury Repair. ACS Appl Mater Interfaces. 2018 May 30;10(21):17742–55.

12. Liu Y, Yu H, Yu P, Peng P, Li C, Xiang Z, et al. Gelatin methacryloyl hydrogel scaffold loaded with activated Schwann cells attenuates apoptosis and promotes functional recovery following spinal cord injury. Exp Ther Med. 2023 Apr;25(4):144.

13. Yan R, Chen S, Wang B, Liu C, Chen X. Magnetic field-oriented conductive decellularized extracellular matrix hydrogel synergizes with electrical stimulation to promote spinal cord injury repair and electrophysiological function restoration. Biomater Adv. 2024 Dec 30;169:214169.

14. Bai J, Liu G, Gao Y, Zhang X, Niu G, Zhang H. Co-culturing neural and bone mesenchymal stem cells in photosensitive hydrogel enhances spinal cord injury repair. Front Bioeng Biotechnol. 2024 Dec 16;12:1431420.

15. Wang H, Tang Q, Lu Y, Chen C, Zhao Y-L, Xu T, et al. Berberine-loaded MSC-derived sEVs encapsulated in injectable GelMA hydrogel for spinal cord injury repair. Int J Pharm. 2023 Aug 25;643:123283.

16. Walsh CM, Wychowaniec JK, Costello L, Brougham DF, Dooley D. An in vitro and ex vivo analysis of the potential of gelma hydrogels as a therapeutic platform for preclinical spinal cord injury. Adv Healthc Mater. 2023 Apr 28;e2300951.

17. Wang Q, Ge L, Guo J, Zhang H, Chen T, Lian F, et al. Acid Neutralization by Composite Lysine Nanoparticles for Spinal Cord Injury Recovery through Mitigating Mitochondrial Dysfunction. ACS Biomater Sci Eng. 2024 Jul 8;10(7):4480–95.

18. Novais GB, Dos Santos S, Santana RJR, Filho RNP, Cunha JLS, Lima BS, et al. Development of a new formulation based on in situ photopolymerized polymer for the treatment of spinal cord injury. Polymers (Basel). 2021 Dec 7;13(24).

19. Li C, Qin T, Zhao J, He R, Wen H, Duan C, et al. Bone Marrow Mesenchymal Stem Cell-Derived Exosome-Educated Macrophages Promote Functional Healing After Spinal Cord Injury. Front Cell Neurosci. 2021 Sep 28;15:725573.

20. Gao Y, Wang K, Wu S, Wu J, Zhang J, Li J, et al. Injectable and photocurable gene scaffold facilitates efficient repair of spinal cord injury. ACS Appl Mater Interfaces. 2024 Jan 7;

21. Morgado PI, Palacios M, Larrain J. In situ injectable hydrogels for spinal cord regeneration: advances from the last 10 years. Biomed Phys Eng Express. 2019 Nov 25;6(1):012002.

22. Piantino J, Burdick JA, Goldberg D, Langer R, Benowitz LI. An injectable, biodegradable hydrogel for trophic factor delivery enhances axonal rewiring and improves performance after spinal cord injury. Exp Neurol. 2006 Oct;201(2):359–67.

23. Omidinia-Anarkoli A, Boesveld S, Tuvshindorj U, Rose JC, Haraszti T, De Laporte L. An Injectable Hybrid Hydrogel with Oriented Short Fibers Induces Unidirectional Growth of Functional Nerve Cells. Small. 2017 Sep;13(36).

24. Parhi R. Cross-Linked Hydrogel for Pharmaceutical Applications: A Review. Adv Pharm Bull. 2017 Dec 31;7(4):515–30.

25. Sandell JL, Zhu TC. A review of in-vivo optical properties of human tissues and its impact on PDT. J Biophotonics. 2011 Nov;4(11–12):773–87.

26. Wang LV, Wu H-I. Biomedical optics: principles and imaging. Hoboken, NJ, USA: John Wiley & Sons, Inc.; 2009.

27. Gąsecka A, Bahdine M, Lapointe N, Rioux V, Perez-Sanchez J, Bonin RP, et al. Light distribution properties in spinal cord for optogenetic stimulation (Conference Presentation). In: Madsen SJ, Yang VXD, Jansen ED, Luo Q, Mohanty SK, Thakor NV, editors. Clinical and Translational Neurophotonics; Neural Imaging and Sensing; and Optogenetics and Optical Manipulation. SPIE; 2016. p. 96902N.

28. DePaoli D, Gasecka A, Bahdine M, Deschenes JM, Goetz L, Perez-Sanchez J, et al. Anisotropic light scattering from myelinated axons in the spinal cord. Neurophotonics. 2020 Jan;7(1):015011.

29. Li H, Zhang C, Feng X. Monte Carlo simulation of light scattering in tissue for the design of skin-like optical devices. Biomed Opt Express. 2019 Feb 1;10(2):868–78.

30. Zhu C, Liu Q. Review of Monte Carlo modeling of light transport in tissues. J Biomed Opt. 2013 May;18(5):50902.

31. Wang H, Wan J, Zhang Z, Hou R. Recent advances on 3D-bioprinted gelatin methacrylate hydrogels for tissue engineering in wound healing: A review of current applications and future prospects. Int Wound J. 2024 Apr;21(4):e14533.

32. Licht C, Rose JC, Anarkoli AO, Blondel D, Roccio M, Haraszti T, et al. Synthetic 3D PEG-Anisogel Tailored with Fibronectin Fragments Induce Aligned Nerve Extension. Biomacromolecules. 2019 Nov 11;20(11):4075–87.

33. Yang K-H, Lindberg G, Soliman B, Lim K, Woodfield T, Narayan RJ. Effect of Photoinitiator on Precursory Stability and Curing Depth of Thiol-Ene Clickable Gelatin. Polymers (Basel). 2021 Jun 5;13(11).

34. Pickering JW, Prahl SA, van Wieringen N, Beek JF, Sterenborg HJ, van Gemert MJ. Double-integrating-sphere system for measuring the optical properties of tissue. Appl Opt. 1993 Feb 1;32(4):399–410.

35. Prahl SA, van Gemert MJ, Welch AJ. Determining the optical properties of turbid mediaby using the adding-doubling method. Appl Opt. 1993 Feb 1;32(4):559–68.

36. S. A. Prahl. Inverse Adding-Doubling Software [Internet]. 2024 [cited 2025 Jan 26]. Available from: https://omlc.org/software/iad/

37. Vogt MR, Hahn H, Holst H, Winter M, Schinke C, Kontges M, et al. Measurement of the Optical Constants of Soda-Lime Glasses in Dependence of Iron Content and Modeling of Iron-Related Power Losses in Crystalline Si Solar Cell Modules. IEEE J Photovoltaics. 2016 Jan;6(1):111–8.

38. Daimon M, Masumura A. Measurement of the refractive index of distilled water from the near-infrared region to the ultraviolet region. Appl Opt. 2007 Jun 20;46(18):3811–20.

39. Jacques SL. Optical properties of biological tissues: a review. Phys Med Biol. 2013 Jun 7;58(11):R37–61.

40. Salomatina E, Jiang B, Novak J, Yaroslavsky AN. Optical properties of normal and cancerous human skin in the visible and near-infrared spectral range. J Biomed Opt. 2006 Dec;11(6):064026.

41. Nickell S, Hermann M, Essenpreis M, Farrell TJ, Krämer U, Patterson MS. Anisotropy of light propagation in human skin. Phys Med Biol. 2000 Oct;45(10):2873–86.

42. Meinke M, Müller G, Helfmann J, Friebel M. Optical properties of platelets and blood plasma and their influence on the optical behavior of whole blood in the visible to near infrared wavelength range. J Biomed Opt. 2007 Feb;12(1):014024.

43. Madelin G, Kline R, Walvick R, Regatte RR. A method for estimating intracellular sodium concentration and extracellular volume fraction in brain in vivo using sodium magnetic resonance imaging. Sci Rep. 2014 Apr 23;4:4763.

44. Leenders KL, Perani D, Lammertsma AA, Heather JD, Buckingham P, Healy MJ, et al. Cerebral blood flow, blood volume and oxygen utilization. Normal values and effect of age. Brain. 1990 Feb;113 (Pt 1):27–47.

45. O’Brien JS, Sampson EL. Lipid composition of the normal human brain: gray matter, white matter, and myelin. J Lipid Res. 1965 Oct;6(4):537–44.

46. Yaroslavsky AN, Schulze PC, Yaroslavsky IV, Schober R, Ulrich F, Schwarzmaier HJ. Optical properties of selected native and coagulated human brain tissues in vitro in the visible and near infrared spectral range. Phys Med Biol. 2002 Jun 21;47(12):2059–73.

47. Sarid H, Abookasis D. Extraction of the anisotropy factor and refractive index of biological tissue in the near-infrared region from diffusion approximation in the spatial frequency domain. Opt Commun. 2022 Apr;508:127749.

48. Leenders KL, Perani D, Lammertsma AA, Heather JD, Buckingham P, Jones T, et al. Cerebral blood flow, blood volume and oxygen utilization. Brain. 1990;113(1):27–47.

49. S. Jacques, T. Li, S. Prahl,. mcxyz.c [Internet]. 2019 [cited 2025 Jan 20]. Available from: https://omlc.org/software/mc/mcxyz/

50. Toossi A, Bergin B, Marefatallah M, Parhizi B, Tyreman N, Everaert DG, et al. Comparative neuroanatomy of the lumbosacral spinal cord of the rat, cat, pig, monkey, and human. Sci Rep. 2021 Jan 21;11(1):1955.

51. Park H, Guo X, Temenoff JS, Tabata Y, Caplan AI, Kasper FK, et al. Effect of swelling ratio of injectable hydrogel composites on chondrogenic differentiation of encapsulated rabbit marrow mesenchymal stem cells in vitro. Biomacromolecules. 2009 Mar 9;10(3):541–6.

52. Advanced BioMatrix - Ruthenium Absorbance Spectrum [Internet]. [cited 2025 Jan 31]. Available from: https://advancedbiomatrix.com/ruthenium-absorbance-spectrum.html

